# Age structuring and spatial heterogeneity in prion protein gene (*PRNP*) polymorphism in white-tailed deer

**DOI:** 10.1101/2020.07.15.205039

**Authors:** Tyler K. Chafin, Marlis R. Douglas, Bradley T. Martin, Zachery D. Zbinden, Christopher R. Middaugh, Jennifer R. Ballard, M. Cory Gray, Don White, Michael E. Douglas

## Abstract

Chronic-wasting disease (CWD) is a prion-derived fatal neurodegenerative disease that has affected wild cervid populations on a global scale. Susceptibility has been linked unambiguously to several amino acid variants within the prion protein gene (*PRNP*). Quantifying their distribution across landscapes can provide critical information for agencies attempting to adaptively manage CWD. Here we attempt to further define management implications of *PRNP* polymorphism by quantifying the contemporary geographic distribution (i.e., phylogeography) of *PRNP* variants in hunter-harvested white-tailed deer (WTD; *Odocoileus virginianus*, N=1433) distributed across Arkansas (USA), including a focal spot for CWD since detection of the disease in February 2016. Of these, *PRNP* variants associated with the well-characterized 96S non-synonymous substitution showed a significant increase in relative frequency among older CWD-positive cohorts. We interpreted this pattern as reflective of a longer life expectancy for 96S genotypes in a CWD-endemic region, suggesting either decreased probabilities of infection or reduced disease progression. Other variants showing statistical signatures of potential increased susceptibility, however, seemingly do so as an artefact of population structure. We also showed marked heterogeneity across the landscape in the prevalence of ‘reduced susceptibility’ genotypes. This may indicate, in turn, that differences in disease susceptibility among WTD in Arkansas are an innate, population-level characteristic that is detectable through phylogeographic analysis.

## Introduction

Chronic wasting disease (CWD) is a fatal neurodegenerative disorder that affects white-tailed deer (WTD; *Odocoileus virginianus*) and related cervids^1,2^, with severe impacts on native wildlife, that also reverberate economically for recreational hunting and ancillary commercial enterprises^3,4^. Most CWD eradication efforts have proven unsuccessful thus far^5^, leading to its continued spread and increased prevalence^6^. Given this, wildlife managers in many jurisdictions are responding by shifting long-term goals away from eradication and instead towards suppression, containment, and mitigation^7,8^.

Several factors have impeded the eradication of CWD, including aspects of life history in both host and agent, as well as limited knowledge with regards to how these interact with environment to define CWD epidemiology. Pathogenicity and transmission, for example, occur via the structural transformation of a naturally occurring cellular prion protein (*PrP*^C^) into a misfolded “pathogenic” isoform (*PrP*^SC^)^9^. The efficiency with which this occurs, coupled with an extensive incubation period^10^, serve to confound proactive surveillance and management. Additionally, both vertical^11,12^ and horizontal transmission^13,14^ are seemingly involved, with prion persistence well established within “environmental reservoirs”^15–17^. As a result, surveillance and monitoring are being increasingly used by state agencies to inform harvest and selective-removal based management strategies to suppress the disease where it cannot be eradicated^18,19^. This is complicated particularly due to a potential for long-distance host dispersal^20–22^, and prion ‘shedding’ by individuals with subclinical infections^23,24^.

Environmental factors that act to reduce disease spread are of particular interest. For example, the capacity of soil as a reservoir for extra-corporeal prion persistence may hinge upon its composition^25^. Likewise, environmental factors that enhance the potential for WTD movements also may modulate CWD transmission among herds^26,27^. Considerable effort to date has focused on characterizing intrinsic susceptibility, especially with regard to the quantification of genetic polymorphisms that encode *PrP*^C^ (*PRNP*)^28–31^. One consistent approach is to identify those *PRNP* gene variants that are associated with reduced CWD susceptibility^32,33^. The amino acid composition of the resulting protein is thus assumed to impact disease progression^34^, although the exact mechanism remains unclear.

Consistent among those variants reported to be associated with reduced CWD susceptibility is a non-synonymous mutation corresponding to an amino acid substitution at position 96 (i.e., from glycine to serine; hereafter 96G versus 96S). Inoculation studies employing this mutation have successfully delayed the progression of CWD in both WTD and proxies^13,35^. Our primary interest is to characterize if (and how) these *PRNP* alleles vary spatially^30,36^, as one landscape-level axis that depicts a “resistance” to CWD spread. We provide such an analysis by employing WTD sampled state-wide as a basis for the phylogeography of the *PRNP* gene (i.e., the geographic distribution of individuals associated with a gene genealogy^37^). We then examine spatial and age-structured patterns within this phylogeographic framework to ask several questions regarding the role of *PRNP* variants on population dynamics in CWD-endemic areas: Do ‘reduced susceptibility’ variants have an effect on survivorship (e.g., as one might expect if *PRNP* polymorphisms do indeed drive susceptibility)? If so, does this leave a *detectable* signature of biased fitness (e.g., as a result of increased survivability and thus potential reproductive output)? What are the impacts of *PRNP* polymorphism on population demographics? We address these questions at two spatial scales: 1) within a dense sampling of the CWD-focal area (Newton County, Arkansas) from which prevalence, and, presumably, measurable impacts on population demography are highest; and 2) state-wide, where sampling densities were lower, and in many areas within which CWD has not yet been detected (though we note that lack of detection does not mean lack of occurrence). The county-level spatial scale allowed us to examine demographic impacts within a recently detected outbreak in a novel area (Newton County), wherein dense sampling and high prevalence allow sufficient sampling for testing hypotheses of age structuring and selection on *PRNP* polymorphisms, while the state-level spatial scale allowed broad-scale analyses of heterogeneity and phylogeographic structuring. The combination of these two spatial scales allows for superimposing differential susceptibilities and fitness across a CWD-absent landscape which could in turn facilitate the creation of management scenarios to project and potentially mitigate disease spread.

## Results

### Data generation

From 2016-2019, ear and tongue tissue samples were collected from 1,720 harvested WTD across 75 counties in Arkansas (Supplemental Table 1; Figure 1). Subsequently, tissue samples from 1,460 WTD were amplified and sequenced, yielding ∼800 nucleotides of the *PRNP* gene and *PRNP*^*psg*^ pseudogene. From these data we obtained 1,433 sequences, of which 316 were obtained from Newton County, the CWD focal area. Sequences were trimmed to 720 unambiguously scored nucleotides, with 11 sites found to be polymorphic (Table 1). Three previously reported polymorphic sites (i.e., nt285, nt299 and nt372; Brandt et al., 2015, 2018) were found to be invariable in our data, whereas one additional site had a novel synonymous substitution (nt499, A/C). Three sites (i.e., nt286, nt367 and nt676) also reflected non-synonymous substitutions, corresponding to amino acid substitutions 96S, 122T and 255K, respectively.

**Figure 1:**
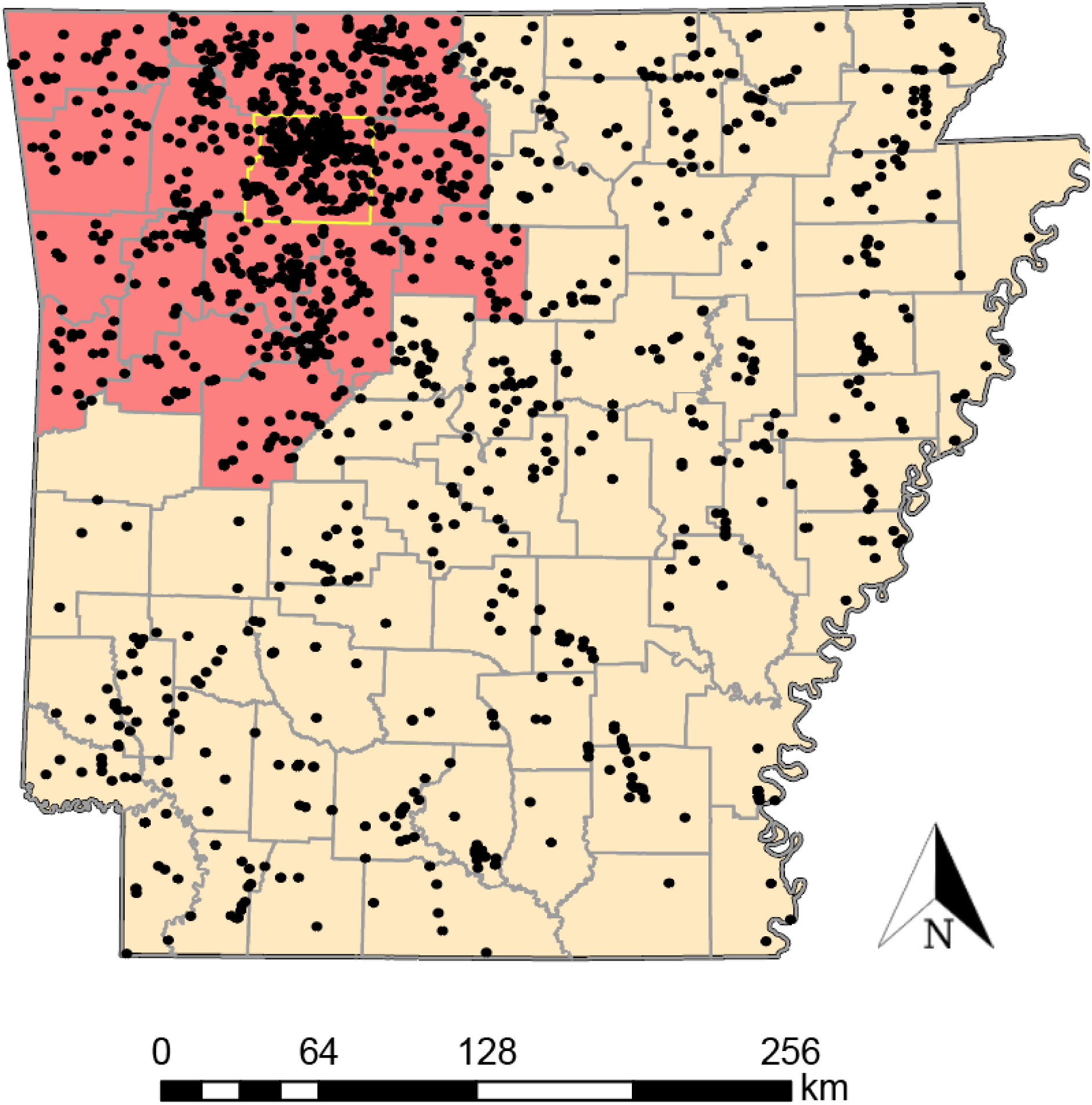
Sampling locations for white-tailed deer tissues from Arkansas evaluated in this study. The red shaded area indicates the 16 counties included in the 2019 Chronic Wasting Disease Management Zone (CWDMZ), with a yellow boundary surrounding a focal area encompassing Newton County. Black dots represent collection localities for each individual tissue. Note that the boundaries of the CWDMZ have since expanded.

**Table 1:**
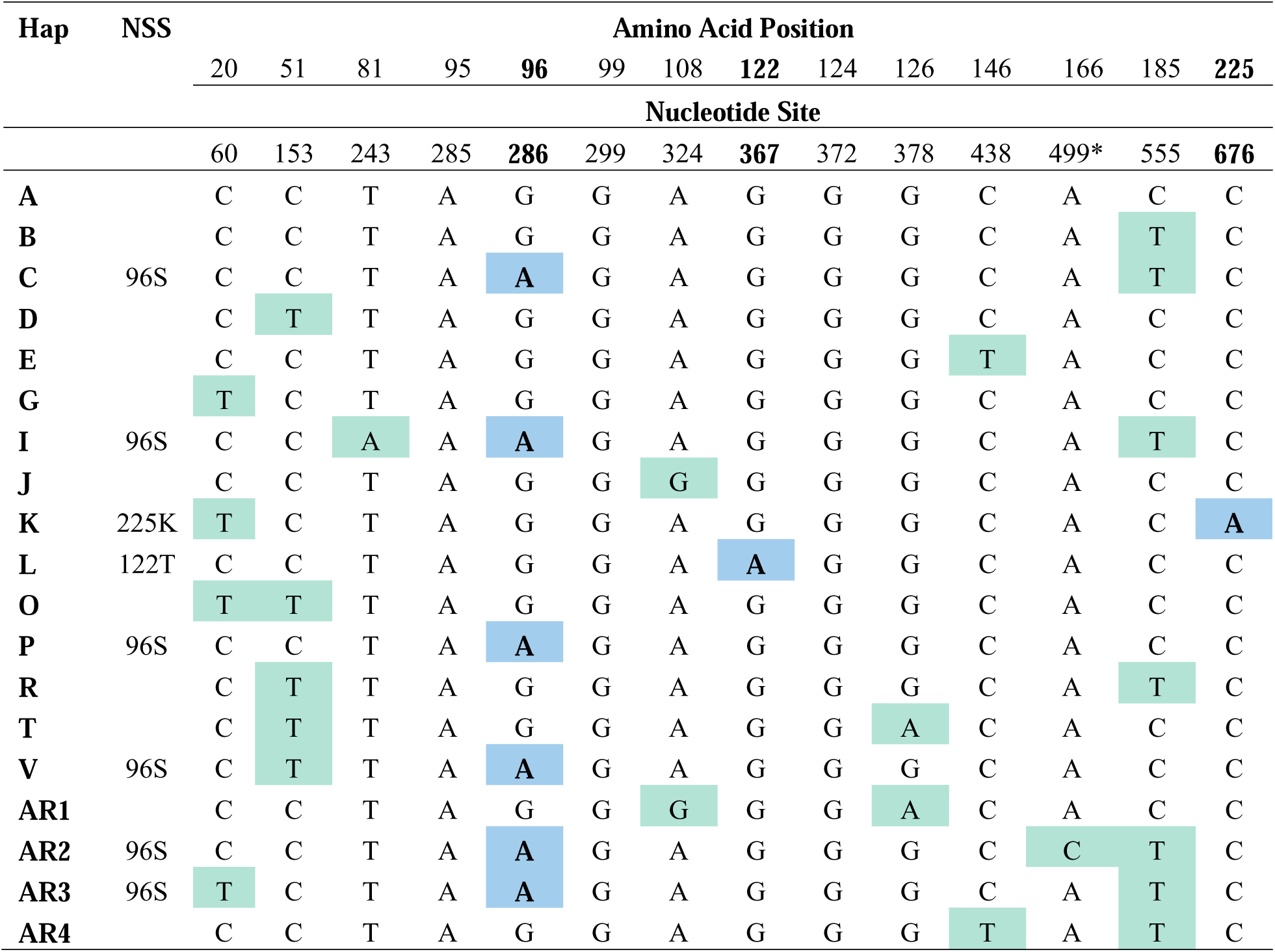
*PRNP* haplotypes tabulated from 1,433 white-tailed deer tissues collected from 75 counties in Arkansas (2016-2019). Haplotypes (Hap) are identified by letter (A-V) following Brandt et al. (2015, 2018). Haplotypes denoted as AR1-4 are previously unreported. Mutations that differ from haplotype A are shaded, with green indicating synonymous substitutions (no amino acid change) and blue as non-synonymous (amino acid changed in protein; NSS). Also listed are amino acid position and nucleotide site (Brandt et al. 2015, 2018).

Haplotype phasing resulted in 20 distinct alleles (Table 1; Figure 2), 4 of which are novel, and 16 were previously documented in other states^29–31^. Two Arkansas haplotypes (AR 2 and AR 3) share the synonymous substitution 96S (nt286/A) associated with reduced CWD susceptibility (i.e., Haplotype C^30^). Three others (I, P and V) also share the 96S amino acid alteration (Table 1; Figure 2). However, all five were at low frequencies (<1%, except Haplotype I at 1.47%; Table 2), and thus were excluded from tests of CWD association.

**Table 2:**
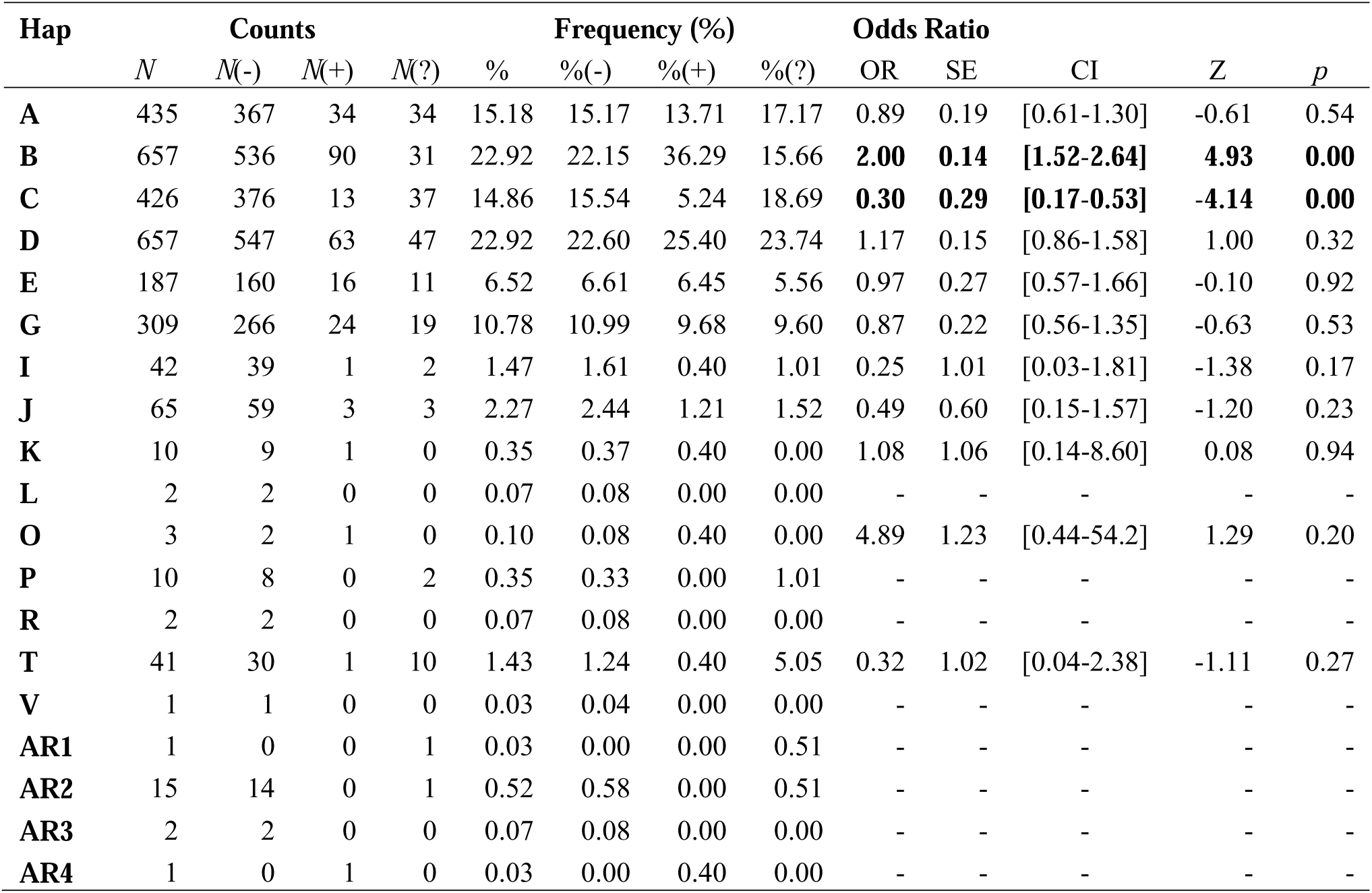
*PRNP* haplotype frequencies and odds-ratios, as associated with CWD status of white-tailed deer in Arkansas, 2016-2019. Haplotypes were derived from unphased sequences representing 720 nucleotides of the *PRNP* gene. Haplotypes (Hap) are identified by letter (A-V) following Brandt et al. (2015, 2018). Haplotypes denoted AR1-4 are previously unreported. Listed are total numbers (N) and relative frequency (%) and associated values for deer that were CWD-negative (-), CWD-positive (+) or untested (?). Odds Ratio (OR) reflects relative representation of a haplotype in CWD-positive deer; OR>1 indicated over-representation, OR<1 under-representation. SE= standard error, CI= 95% confidence interval, Z= OR Z-score and *p*= OR *p*-value. Values in bold are significant.

**Figure 2:**
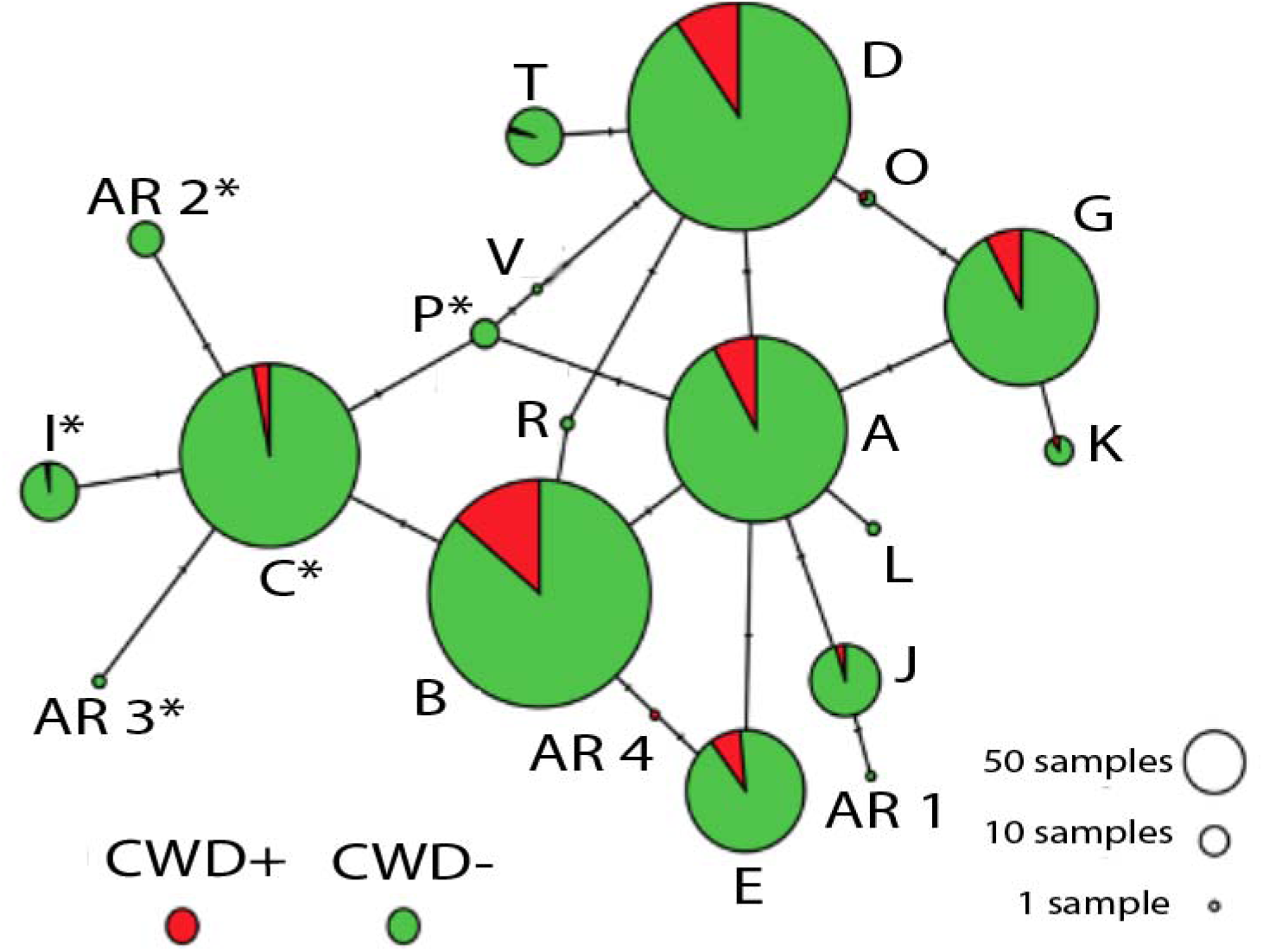
Haplotype network showing relationship of prion gene variants (*PRNP*) detected across 1,433 white-tailed deer collected from 75 counties in Arkansas (2016-2019). Data are based on sequence analysis of 720 nucleotides. Circles represent 20 haplotypes (=alleles) with size reflecting frequency of occurrence in entire data set (Table 3), and tick marks representing number of mutations (=nucleotide substitutions) distinguishing one from another (Table 2). Color codes reflect relative frequency among CWD-positive (red) and CWD-negative/ undetected (green). Letters correspond to haplotype names (per Brandt et al. 2015), with haplotypes unique to Arkansas indicated with numbers (AR#). Haplotypes sharing the 96S mutation are indicated with (*).

Targeted amplification and sequencing of the *PRNP*^*PSG*^ pseudogene was successful in 30% of our samples (443 of 1,459). Our comparison of the 443 *PRNP* and *PRNP*^*PSG*^ haplotypes indicated that variability at site nt413 did not represent a biological variant of the *PRNP* gene but was instead a non-targeted amplification of the *PRNP*^*PSG*^ pseudogene.

### PRNP haplotype frequencies

Haplotype frequencies (Tables 2 and S2) differed slightly from those reported in other states (e.g., Wisconsin and Illinois^30,31^). The four most frequent haplotypes in Illinois and Wisconsin (i.e., <10%), were also common among Arkansas samples (Figure 3; Haplotypes A-D). We also found that Haplotype A, most common in Illinois/Wisconsin at 30%, occurred in only 15% of our samples (Figure 3; Table 2). Haplotypes B and D were instead most common in our data (each at ∼23%). Haplotype C, associated with reduced CWD susceptibility, was detected at a frequency of 15%, quite similar to that in Illinois/Wisconsin (17%). Two additional haplotypes (E and G) also occurred at high frequencies in Arkansas (7% and 11%, respectively), whereas they were found at <5% in the Illinois/Wisconsin^30^. Out of the 16 rare haplotypes with previously reported frequencies ≤1% (Haplotypes K-Z^30^), seven were also observed at low frequencies in Arkansas (i.e., Haplotypes K, L, O, P, R, T, and V). Haplotype frequencies also differed across counties (Supplemental Table 1), although we note that sampling effort was uneven across the state.

**Figure 3:**
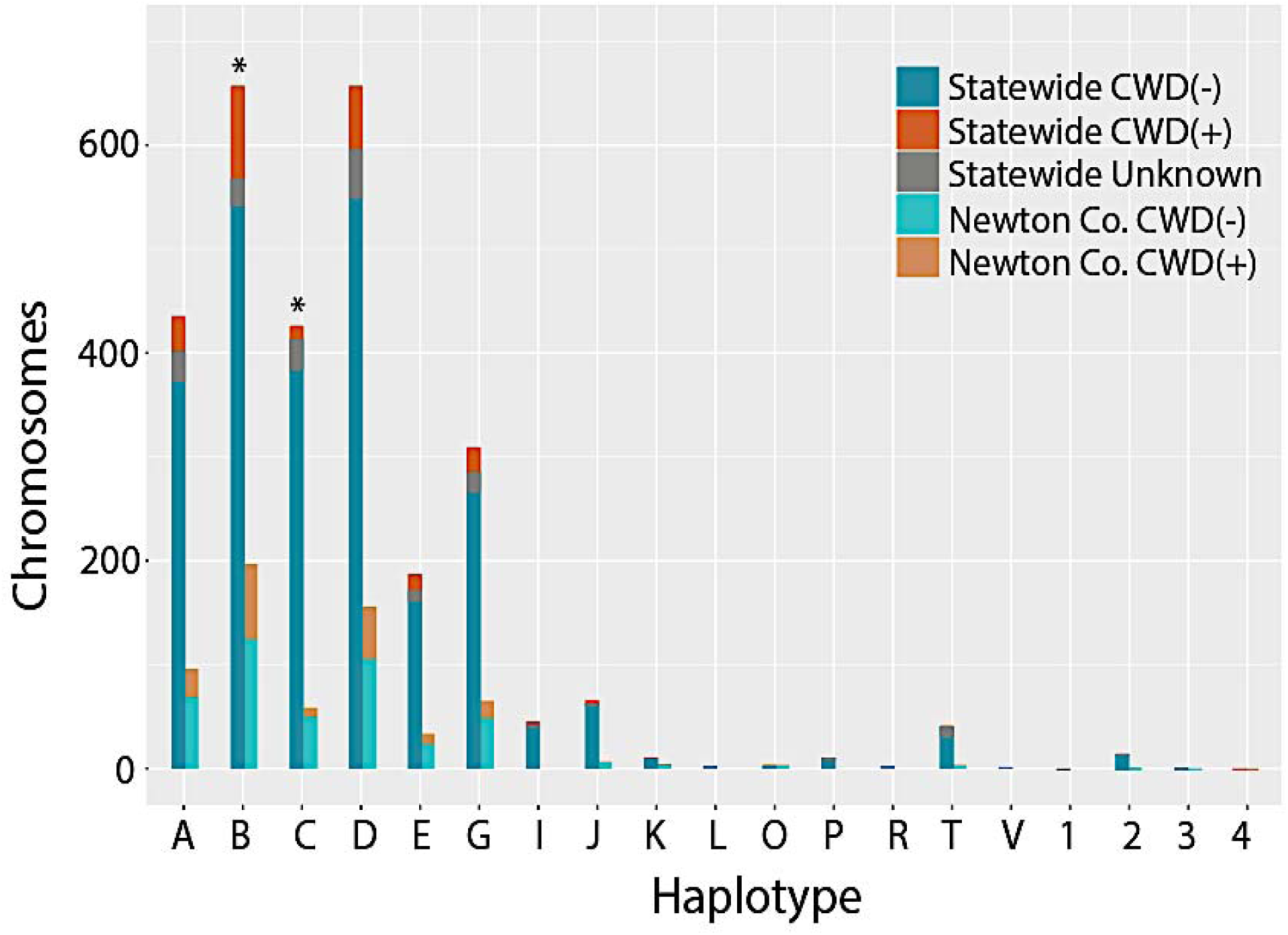
Frequency distribution of 2,866 *PRNP* haplotypes detected in white-tailed deer collected in Arkansas, 2016-2019. Haplotypes were determined by phasing individual genotypes derived from sequencing 1,433 deer across 720 bp of the *PRNP* gene. Letters (A through V) refer to haplotypes identified in Brandt et al. (2015), whereas numbers (1-4) are haplotypes unique to Arkansas, and thus previously unreported. Frequencies are plotted for all 1,433 samples (=statewide) and a subset of 314 samples from Newton County (N=628 chromosomes). Color codes reflect frequency among CWD-positive (CWD+) and CWD-negative (CWD-) samples; unknown indicates samples that were not tested for CWD.

### Evidence for disease-mediated selection on PRNP variants

Haplotypes B and C did not occur in the same frequency within CWD-positive (*N* = 432) and CWD-negative deer (*N* = 196; Table 2; Figure 3). Haplotype B was over-represented within CWD-positive deer, both state-wide (OR = 2.00, *p* = 0.000001), and in Newton County (OR = 1.43, *p* = 0.033). Haplotype C was under-represented within CWD-positive deer, both state-wide (OR = 0.30, *p* = 0.00003) and Newton County (OR = 0.42, *p* = 0.015).

If Haplotypes B and C influence CWD susceptibility, then their relative frequency of occurrence should vary among deer age-classes, reflecting a biased probability of reaching older age classes. We restricted our analyses to Newton County to ensure only deer from CWD-endemic locations were considered. We found the frequency of Haplotype C significantly more common in older deer, both when measured as relative allele frequency (*p*=0.036; *R*^2^=0.635; Figure 4), and as an age-partitioned odds ratio (*p*=0.043; *R*^2^=0.601; Figure 5). The increase in Haplotype C remained substantial even when we considered only the relative frequency of each haplotype among CWD-positive deer from Newton County (i.e., 4% of yearling/ fawn haplotypes, and 17% of those from individuals older than 5; Supplemental Table 3). Among CWD-positive deer state-wide, Haplotype C was recorded at 5.24% of all sampled haplotypes (Table 2). By contrast, Haplotype B showed neither a discernible relationship regarding CWD status, nor relative frequencies across age classes (Figure 5; Supplemental Table 3).

**Figure 4:**
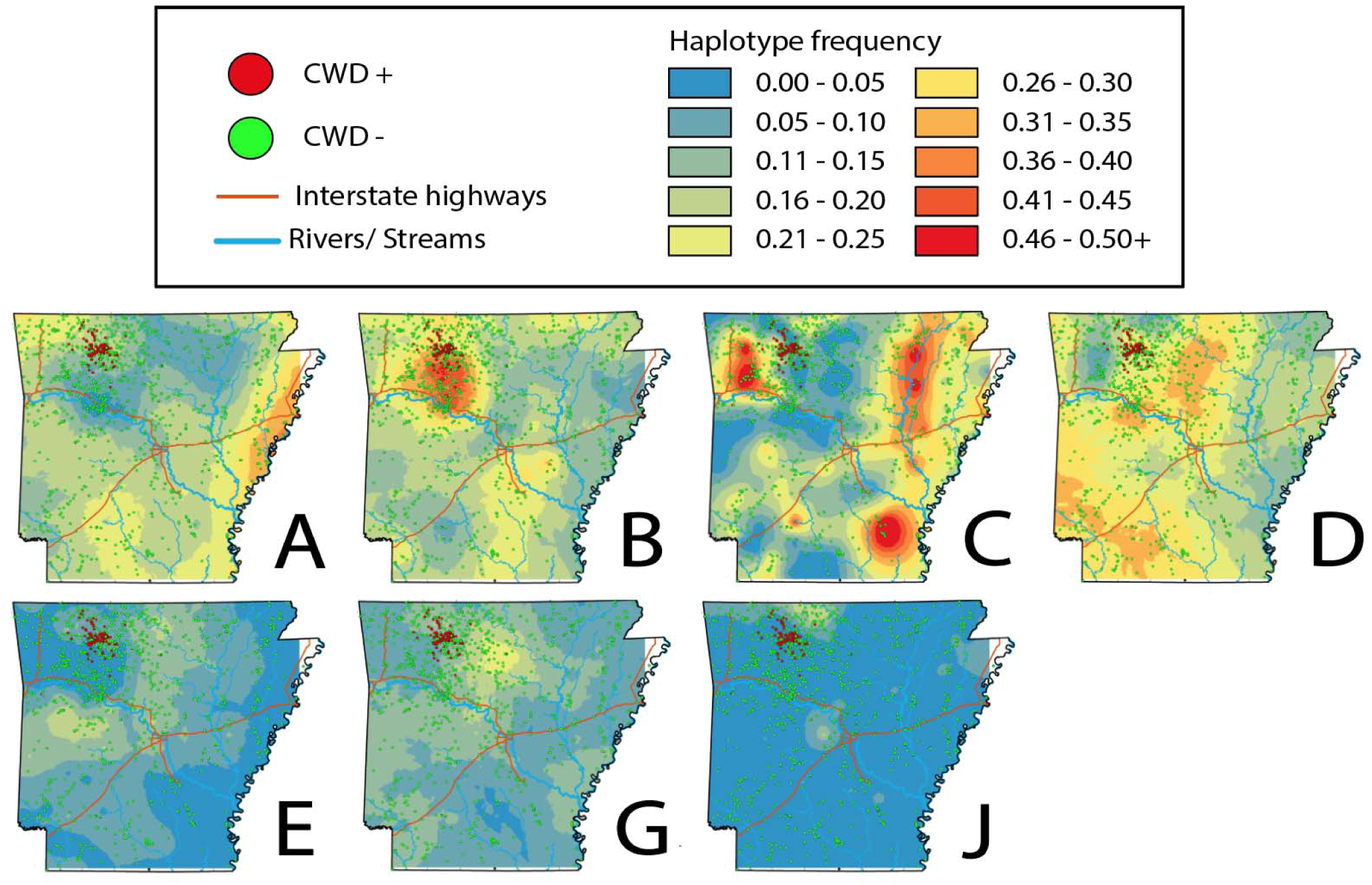
Topographies that represent interpolated haplotype frequencies for the *PRNP* gene in Arkansas. Frequency is depicted by color, with blue reflecting low occurrence (0-5%) whereas red indicating 46-50+% of haplotypes were of this type.

**Figure 5:**
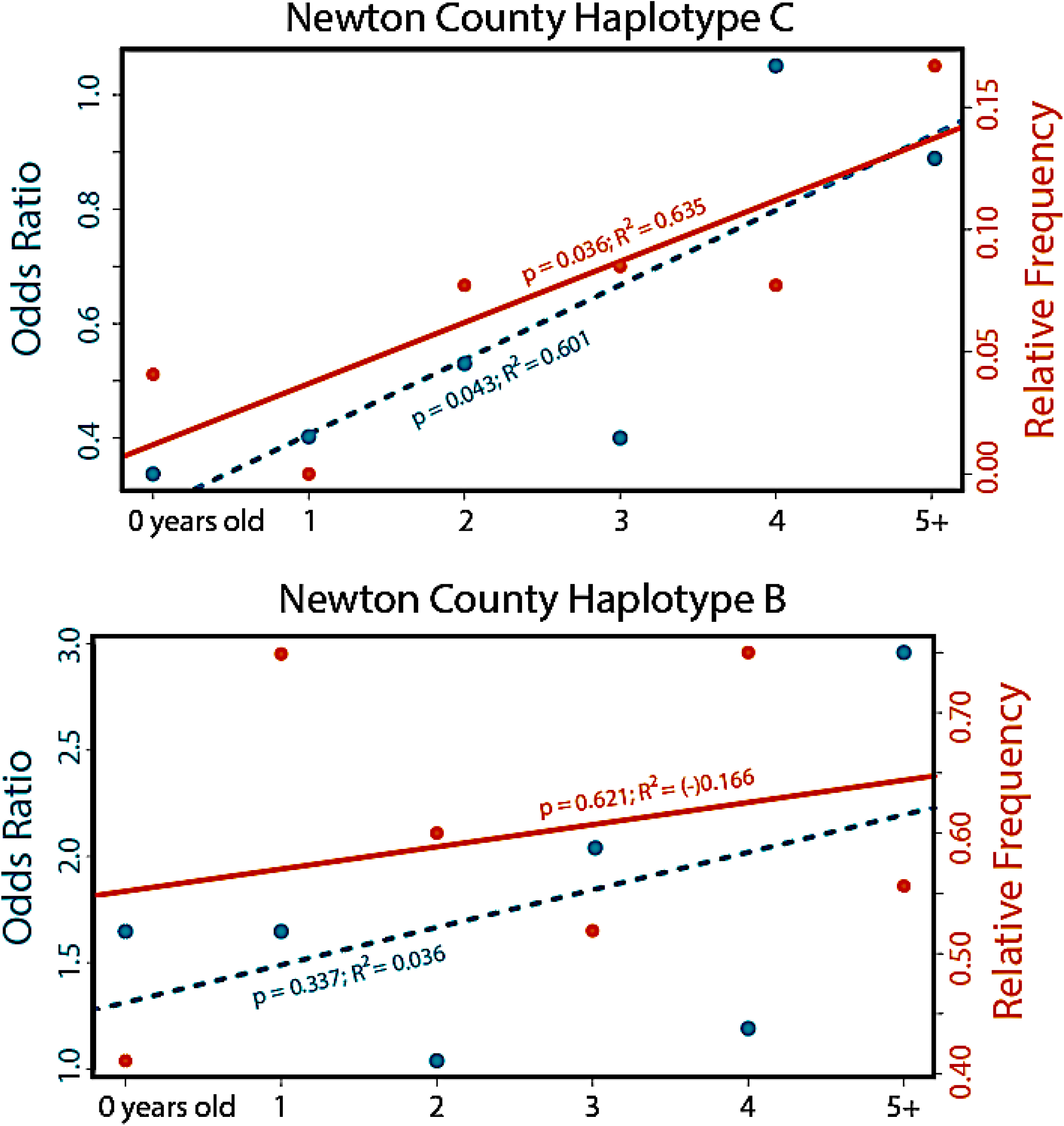
Relative frequency and odds ratio for two haplotypes of the prion gene *PRNP* haplotypes detected in white-tailed deer age cohorts (<1 year to 5+ years) sampled in Arkansas from 2016-2019. Prion gene variant Haplotype C (top panel) has been associated with reduced susceptibility to CWD, whereas Haplotype B (lower panel) has been associated with higher susceptibility (Brandt et al. 2018). Data are based on phased haplotypes derived from 720 nucleotides of the *PRNP* gene sequenced across 1,433 deer.

Given the presence of age-structuring in Haplotype C, we then hypothesized a fitness differential associated with polymorphism at AA position 96. We first examined phylogenetic conservation and stability of non-synonymous mutations in Arkansas (Table 1) and predicted subsequent effects on tertiary protein structure. We found the 96S substitution to be the least conserved of the three amino acid positions, with none of the observed substitutions significantly altering tertiary shape (Supplemental Table 3). We thus hypothesized that positive selection on 96S was driving the observed patterns. As a test, we treated cohorts in Newton County as a time-series and computed the magnitude of selection required to generate the observed positive growth in frequency with increasing age (Figure 5). Our results yielded a selection coefficient (*s*) of 0.1215. We note that numerous caveats are associated with population genetic assumptions behind this calculation, as well as of the interpretation of selection coefficients^38^, the result seemingly supports our observations.

## Discussion

Our diagnosis of variability in the *PRNP* gene across 1,433 WTD collected from 75 counties in Arkansas was comparable to that found in other states (Tables 1-2). Of the 20 haplotypes we detected, 16 were previously identified in other states, with those alleles at higher frequencies in Arkansas also being most common elsewhere^30^. Four novel variants were found at low frequencies (Table 2; Figure 2-3), with paralog artefacts as a source of this variation eliminated due to our sequencing of the pseudogene. Haplotype C, characterized by the 96S substitution, showed a significantly biased representation among CWD-positive deer, being under-represented in younger deer and over-represented in older CWD-positive deer (Figure 5). This suggests that 96S-individuals tend to live longer following CWD-infection than those with alternate genotypes. This could reflect either a reduced likelihood of contracting the disease or a slower disease progression following exposure.

We also found Haplotype B as significantly over-represented in CWD-positive deer (Table 2 and Figure 3) but failed to find a reciprocal impact on life expectancy (Figure 5). Instead, we found Haplotype B to be most extensive within the region where CWD is currently centered (Figure 4). This, in turn, may suggest heterogeneity in Haplotype B frequency represents an artefact of population structure, a well-known confounding variable recognized in trait-association studies^39–42^. We also note that decreased precision of aging of white-tailed deer on the basis of tooth development and wear patterns may also contribute some variability, particularly with regard to haplotype frequencies among older cohorts^43–45^. We also cannot rule out the potential for linked variation as influencing the observed pattern in Haplotype B, however our results do not find any evidence for increased susceptibility of the variant. Using panels of nuclear markers, we suggest further study of patterns of spatial connectivity and population structure in wild WTD^46^ as a means of separating potential spatial and phylogeographic drivers of haplotype frequencies from those driven by disease-mediated selection.

### Management implications of PRNP variation

Structural variants of the *PrP* protein play a role in disease progression^28,34^. However, the exact mechanisms remain poorly understood. The primary variant we have implicated herein (i.e., 96S) as influencing disease susceptibility has been supported as such in laboratory settings. For example, Mathiason et al.^23^ inoculated WTD with prion strains and examined time-to-detection (via saliva) across deer in multiple cohorts. Several infected individuals having the 96S prion gene variant remained undetected at 18 months post-inoculation, although this may represent an insufficient time for the necessary *in vivo* prion protein build-up. Race et al.^35^ similarly inoculated transgenic mice expressing different white-tailed deer 96GG and 96SS *PRNP* genotypes and showed that this delay in disease progression also extended to heterozygotes (96GS genotype).

Despite compelling evidence for an inhibitory mechanism at the 96S allele, its use as a management tool remains unclear. Genetically-guided selective breeding in domestic sheep (*Ovis aries*) has reduced scrapie incidence ^47,48^. In a captive setting such an approach may be viable for WTD^35^, although we note that the degree of protection offered by the 96S genotype to CWD is likely much lower than that seen among the most ‘protective’ genetic variants to scrapie in sheep^47^. However, captive deer maintained at high density and under genetic selection could also drive artificial selection for emergent prion strains with heightened pathogenicity in 96S deer, as well as potentially expanding the host range to include novel species^49^.

The implication of 96S frequency with regard to the spread of CWD within and among herds is likewise uncertain. An important question is whether the potential for an increased incubation period ^50^ associated with 96S could also produce a similar extended period for asymptomatic and subclinical transmission/prion shedding^24,51^. If so, this could increase the probability of transmission by 96S deer, thus promoting its increase within populations (Figure 6). An understanding of the of prion growth kinetics across host genotypes is needed, as well as a more thorough understanding of prion strain evolution^52^.

**Figure 6:**
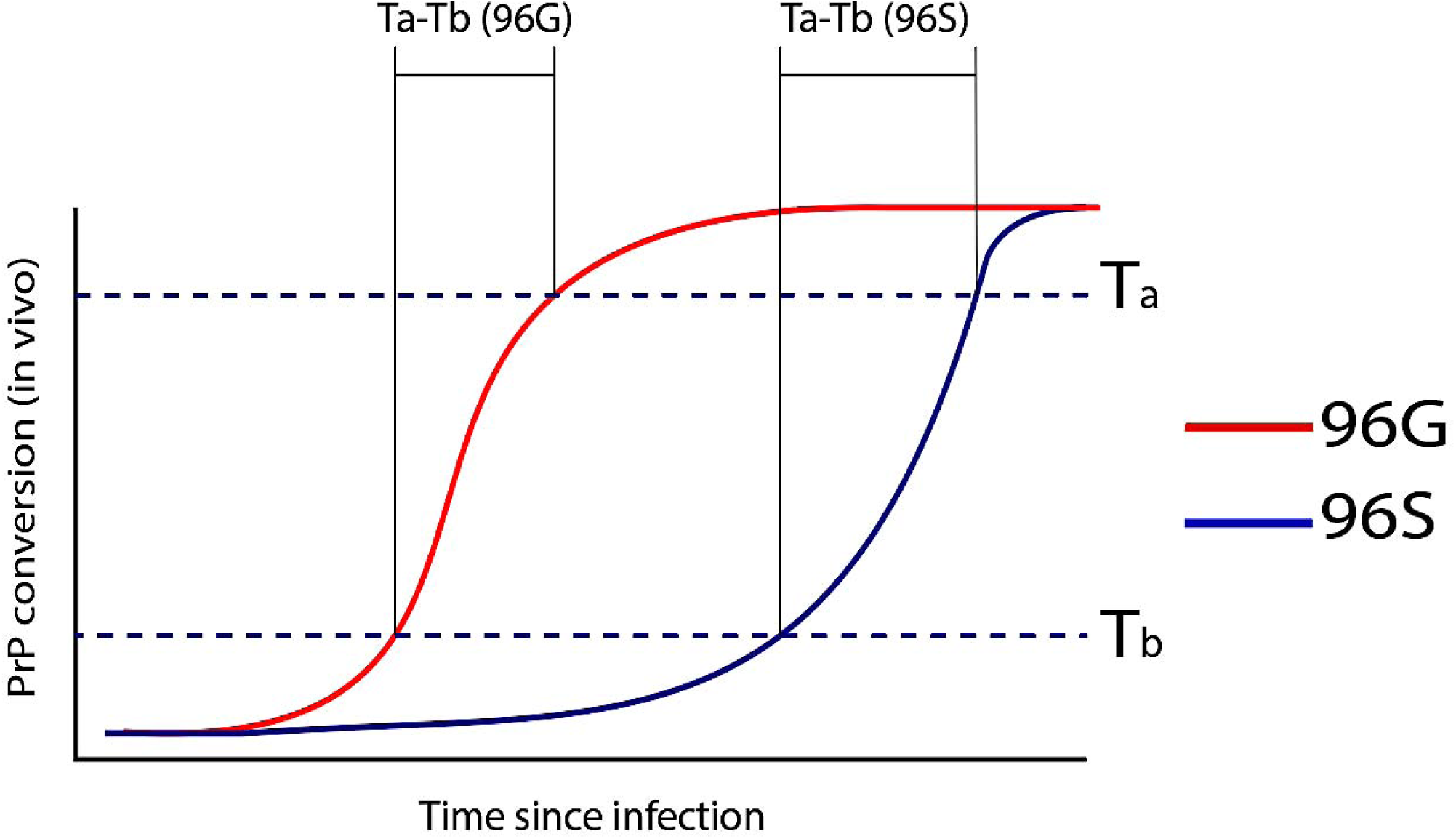
Results of a conceptual model depicting prion conversion rates for *PrP*^C^ variants 96G and 96S. The diagram demonstrates prion conversion of 96G and 96S (reduced susceptibility) prion gene (=*PRNP*) variants. Theoretical prion protein (=*PrP*) mass thresholds were imposed: Ta = time of symptomatic expression; and Tb = theoretical time after which individuals remain asymptomatic. Individual-individual transmission is possible through direct contact (saliva) or shedding (e.g. feces). The time interval for *asymptomatic spread* (e.g., prion shedding from a subclinical individual) extends prior to *PrP* conversion surpassing threshold Tb, but not extending beyond Ta.

## Conclusion

Our results corroborate previous research conducted with WTD in Illinois and Wisconsin: reduced CWD susceptibility in *PRNP* variants associated with the non-synonymous 96S mutation^30,31^. We demonstrated that a common haplotype (Haplotype C) harboring 96S increased in relative frequency when older CWD-positive cohorts were examined.

Management implications of this research requires further epidemiological understanding necessary to predict outcomes (Figure 6). The next step in understanding the current distribution and future spread of CWD in Arkansas requires the characterization of context-specific factors, including 1) population structure as a potential driver of *PRNP* trends, 2) landscape features that modulate deer dispersal, population demography, and density, and 3) increased knowledge of epidemiology, including the interactions among context-specific factors^53^. We thus advocate for a landscape genomic framework for WTD as the next logical step to characterize CWD spread such that fine-scaled deer movement patterns can be more effectively parsed and interpreted^54^.

## Materials and Methods

### CWD surveillance and prevalence in Arkansas

In October 2015, CWD was initially detected in Arkansas in a 2.5-year old female elk (*Cervus canadensis*) legally harvested near Pruitt in Newton County. In February 2016, a CWD-positive female WTD was found dead in Ponca in Newton County. During March 2016, biologists from the Arkansas Game and Fish Commission (AGFC) collected 266 WTD tissue samples within a 50,500 ha focal region. CWD prevalence was 23% and differed by gender (female = 20%, male = 32%). For these surveys, to include the samples collected for molecular work, aging of deer was done by examination of tooth replacement and wear^55^.

Subsequent state-wide monitoring, which included hunter-harvested and road-killed deer, identified CWD-positive individuals outside of the initial focal region. Additional state-wide sampling efforts, in conjunction with hunter harvested and bi-annual surveys, established a state-wide baseline for occurrence of CWD. As of April 2020, 818 WTD and 23 elk have tested positive for CWD. Given the incidence of confirmed CWD-positive deer, AGFC established a CWD Management Zone (CWDMZ) that included counties within a 16km radius of identified positives. At the completion of this study (June 2019), the CWDMZ encompassed 19 counties of northwestern Arkansas (Figure 1).

There has been a concerted effort by the AGFC to proactively manage CWD prevalence and potential disease spread in Arkansas, which included hunting regulations that promote harvest of young male deer and increased harvest of female deer (e.g., removal of antler point restrictions and altered bag limits). Addition restrictions included prohibiting baiting and feeding to reduce grouping behavior in deer and to hinder human-mediated transmission via hunting and subsequent removal of carcasses from with the CWDMZ. The CWDMZ has subsequently been expanded to encompass the known distribution of CWD. During the 2018/2019 deer and elk hunting seasons, 246 additional CWD-positive cervids (241 WTD and five elk) were detected. Moreover, the AGFC has mandated a compulsory testing requirement for harvested elk, and a voluntary test for WTD, facilitated by a state-wide network of deer head drop-off locations.

### DNA Extraction and amplification

Frozen ear or tongue tissue was homogenized with the TissueLyser II (QIAGEN^©^ Corporation, Maryland, USA), with genomic DNA subsequently extracted using the QIAamp Fast DNA Tissue Kit protocol (QIAGEN^©^ Corporation, Maryland, USA). To ascertain presence of high-quality genomic DNA (i.e., molecular weight >20kb), a 5μl aliquot of the DNA extract was separated on a 2% agarose gel and visualized using GelGreen on a blue-light transilluminator (Gel Doc(tm) EZ Imager; Bio-Rad). DNA was then used as template material to amplify a coding section of the *PRNP* gene, following a protocol modified from previous studies^31,32^. For the functional *PRNP* gene, the forward primer (CWD-13) straddles Intron 2 and Exon 3, with the reverse primer (CWD-LA) located 850bp downstream^32^. To ascertain if the polymorphisms were indeed in the functional *PRNP* gene, we tested for the presence of the non-coding *PRNP* pseudogene (*PRNP*^*PSG*^) by using pseudogene primers 223 and 224^33^.

Amplifications for the functional *PRNP* gene and the *PRNP*^*PSG*^ pseudogene were performed in 20μl reactions consisting of 10μl Qiagen HotStart Master Mix (1unit HotStartTaq DNA Polymerase, PCR Buffer with 3mM MgCl_2_, and 400µM of each dNTP), 8μM each of the forward and reverse primer, 7.4μl DNase-free water, and 1μl of template DNA (∼50-100ng). Thermocyling protocols consisted of an initial denaturation step of 15min at 95°C, followed by 10 cycles of 45s denaturation at 95°C, 45s annealing at 57°C, and 75s extension at 72°C, 25 cycles of 30s denaturation at 95°C, 30s annealing at 55°C and 60s extension at 72°C, completed with a final extension step of 5min at 72°C.

If both *PRNP* and *PRNP*^*PSG*^ amplified in a sample, then each was sequenced to identify the true polymorphism in the functional *PRNP* gene. Amplicons were enzymatically purified, sequenced using BigDye v. 3.1 (Applied Biosystem Inc., Forest City, CA) dye-terminator chemistry, and resolved on an ABI 3730XL GeneAnalyzer (University of Illinois Keck Center for Functional and Comparative Genomics). Sequences were manually edited using Sequencher (v 5.4, Gene Codes, Ann Arbor MI) and aligned against a reference database of *PRNP* gene sequences obtained from the NCBI GenBank (Accession # AF156185.1; AY3600089.1; AY3600091.1).

### Haplotype phasing and network construction

Following alignment, sequences were phased to haplotypes (paired nuclear alleles) using the program Phase2^56^, which reconstructs haplotypes using a probabilistic model of linkage disequilibrium. Only haplotypes assigned with >90% posterior probability (N=1,433) were retained. Scripts employed to format inputs and parse haplotype phasing are available at *github*.*com/tkchafin/haploTools*. Haplotypes were then categorized using a published nomenclature^31^, with haplotype frequencies calculated globally, by county, and by CWD status. We constructed a haplotype network to visualize similarity amongst haplotype, using the median-joining algorithm employed by PopArt^57^. Scripts to formulate these input files are found at: *github*.*com/tkchafin/scripts*.

### Analysis of PRNP variants

We first applied spatial interpolation to examine the structure of *PRNP* haplotypes as distributed across the state. Haplotype frequencies were computed by first dividing the state into non-overlapping ‘pseudo-populations’ that contained between 5-10 sampling localities each. This was done because our state-wide sampling process lacked *a priori* information with regard to natural population structure. Our results yielded N=211 polygons (Supplemental Figure 1). We then applied Empirical Bayesian Kriging in ArcMap v10.7.1 (Esri, Inc.) to interpolate our posterior probabilities.

To associate disease with *PRNP* variants, we followed prior studies^30,31^ by computing odds-ratios (OR). We first consider the probability of displaying an outcome (=CWD status) given the presence of a focal haplotype. An OR>1 = an over-representation of the focal haplotype among CWD-positive deer; an OR<1 indicates the focal haplotype is under-represented. We identified and evaluated haplotypes separately, by examining haplotype frequencies among age classes. We did so because a bias in relative representation can be driven by multiple factors such as population structure, which may drive a coincident relationship. Here, we assumed that if a haplotype indeed affects the probability of survival to adulthood in CWD-present regions, presumably by reducing disease risk and/or progression, then it should show a significant change in relative representation in older age groups.

We then computed a selection coefficient (*s*) as a relative fitness value. We first derived counts for each haplotype within each age class, then sampled across age classes as a time-series sample. We then estimated selection by examining changes in allele frequencies across this series, relative to that expected via stochastic change^58^. Finally, we modelled the effects non-synonymous mutations on protein conformation so as to test if they do indeed alter *PrP*^*SC*^ tertiary structure (as opposed to simply being linked to un-sequenced variation). To understand amino acid sequence conservation and possible deleterious effects at each non-synonymous position, we utilized two alignment-based algorithms, PROVEAN^59^ and SIFT^60^. We also predicted protein stability using the support vector machine (SM)-based I-Mutant2.0 tool^61^.

## Acknowledgments

The authors thank hunters and wildlife managers statewide who collected WTD tissue samples. We give particular thanks to A.J. Riggs for helping to coordinate sample collection. We also thank former students and technicians P. McDill and L. Graham for contributions to study design, sample handling and organization, and DNA extraction.

## Disclosure statement

No potential conflict of interest is reported by the authors.

## Funding

This project and the preparation of this publication was funded in part by the Pittman-Robertson Wildlife Restoration Grant Program (Grant #AR-W-F18AF00020) of the U.S. Fish and Wildlife Service through an agreement with the Arkansas Game and Fish Commission. The views and opinions expressed herein are those of the authors and do not necessarily reflect the views or policies of the Arkansas Game and Fish Commission. Product references do not constitute endorsements.

## Supplemental Figures and Tables

**Supplemental Table 1:** Samples per county and frequency of 20 *PRNP* haplotypes detected in 1,433 white-tailed deer collected in 75 counties in Arkansas from 2016-2019. Phased haplotypes were derived from sequence analysis of 720 nucleotides of the *PRNP* gene. Letters indicate haplotypes previously detected in other states, whereas numbers (1-4) indicate haplotypes unique to Arkansas. Variable sites of haplotypes are listed in Table 2. Samples that tested positive for CWD (+) are listed separately for those counties where CWD was detected.

| County | N | Haplotype |  |  |  |  |  |  |  |  |  |  |  |  |  |  |  |  |  |  |
| --- | --- | --- | --- | --- | --- | --- | --- | --- | --- | --- | --- | --- | --- | --- | --- | --- | --- | --- | --- | --- |
|  |  | A | B | C | D | E | G | I | J | K | L | O | P | R | T | V | 1 | 2 | 3 | 4 |
| Arkansas | 11 | 4 | 5 | 5 | 4 | - | 3 | - | - | - | - | - | - | - | 1 | - | - | - | - | - |
| Ashley | 1 | 1 | - | - | - | - | - | - | 1 | - | - | - | - | - | - | - | - | - | - | - |
| Baxter | 16 | 3 | 7 | 3 | 8 | 4 | 6 | - | 1 | - | - | - | - | - | - | - | - | - | - | - |
| Benton | 40 | 25 | 19 | 3 | 16 | 5 | 5 | 1 | 4 | - | - | - | 1 | - | 1 | - | - | - | - | - |
| Boone | 33 | 6 | 24 | 8 | 7 | 5 | 8 | - | 7 | - | 1 | - | - | - | - | - | - | - | - | - |
| Boone(+) | 2 | - | - | - | 3 | - | 1 | - | - | - | - | - | - | - | - | - | - | - | - | - |
| Bradley | 2 | 1 | - | 2 | - | - | - | - | - | - | - | - | 1 | - | - | - | - | - | - | - |
| Carroll | 33 | 18 | 13 | 4 | 11 | 4 | 7 | - | 6 | - | - | - | - | - | 3 | - | - | - | - | - |
| Carroll(+) | 16 | 5 | 12 | 2 | 5 | 3 | 5 | - | - | - | - | - | - | - | - | - | - | - | - | - |
| Calhoun | 14 | 4 | 9 | 1 | 6 | 3 | 2 | - | 1 | - | - | - | 1 | - | 1 | - | - | - | - | - |
| Cleburne | 8 | 2 | 2 | 2 | 4 | 2 | 4 | - | - | - | - | - | - | - | - | - | - | - | - | - |
| Chicot | 4 | 3 | - | 1 | 2 | - | 1 | 1 | - | - | - | - | - | - | - | - | - | - | - | - |
| Clark | 13 | 2 | 5 | 6 | 6 | 1 | 1 | 3 | 1 | - | - | - | 1 | - | - | - | - | - | - | - |
| Conway | 10 | 1 | 4 | 1 | 6 | 3 | 5 | - | - | - | - | - | - | - | - | - | - | - | - | - |
| Columbia | 4 | 1 | 1 | - | 4 | - | 2 | - | - | - | - | - | - | - | - | - | - | - | - | - |
| Craighead | 12 | 5 | 1 | 5 | 7 | 1 | 3 | - | 1 | - | - | - | - | - | 1 | - | - | - | - | - |
| Cross | 9 | 4 | 4 | 3 | 5 | - | 1 | 1 | - | - | - | - | - | - | - | - | - | - | - | - |
| Crittenden | 8 | 5 | 1 | 5 | 4 | - | 1 | - | - | - | - | - | - | - | - | - | - | - | - | - |
| Crawford | 9 | 1 | 3 | 5 | 6 | - | 1 | - | - | 2 | - | - | - | - | - | - | - | - | - | - |
| Clay | 14 | 7 | 4 | 7 | 3 | 1 | 2 | 1 | - | - | - | - | 1 | - | 2 | - | - | - | - | - |
| Dallas | 9 | 6 | 3 | 1 | 3 | 3 | 1 | 1 | - | - | - | - | - | - | - | - | - | - | - | - |
| Drew | 14 | 4 | 2 | 16 | 3 | 1 | 2 | - | - | - | - | - | - | - | - | - | - | - | - | - |
| Desha | 12 | 8 | 4 | 5 | 4 | - | 1 | - | - | - | - | - | - | - | 1 | - | 1 | - | - | - |
| Faulkner | 14 | 5 | 1 | 2 | 11 | 4 | 1 | - | 1 | - | - | - | 1 | 2 | - | - | - | - | - | - |
| Franklin | 33 | 6 | 23 | 29 | 1 | - | 5 | - | - | - | - | - | - | - | - | - | - | 2 | - | - |
| Fulton | 21 | 3 | 8 | 4 | 12 | 4 | 5 | - | 1 | - | - | - | - | - | 5 | - | - | - | - | - |
| Garland | 16 | 6 | 5 | 6 | 5 | 4 | 1 | 3 | - | - | - | - | - | - | - | - | - | 2 | - | - |
| Greene | 12 | 1 | 8 | 7 | - | - | 2 | 2 | 1 | - | - | - | - | - | 3 | - | - | - | - | - |
| Grant | 10 | 4 | 3 | 3 | 7 | - | 2 | - | - | - | - | - | - | - | 1 | - | - | - | - | - |
| Hampstead | 10 | 6 | 3 | - | 5 | 2 | 1 | 3 | - | - | - | - | - | - | - | - | - | - | - | - |
| Howard | 13 | - | 3 | 6 | 11 | 2 | 2 | 1 | - | - | - | - | - | - | 1 | - | - | - | - | - |
| Hot Springs | 5 | 2 | 2 | 2 | 1 | 1 | 2 | - | - | - | - | - | - | - | - | - | - | - | - | - |
| Independence | 10 | - | 3 | 5 | 4 | 1 | 4 | - | - | - | - | - | - | - | 1 | - | - | 2 | - | - |
| Izard | 9 | - | 2 | - | 7 | 6 | 2 | - | - | - | - | - | - | - | - | - | - | 1 | - | - |
| Jackson | 2 | - | - | 3 | 1 | - | - | - | - | - | - | - | - | - | - | - | - | - | - | - |
| Jefferson | 13 | 6 | 7 | 2 | 5 | 1 | 4 | - | 1 | - | - | - | - | - | - | - | - | - | - | - |
| **Johnson** | **45** | 10 | 33 | 19 | 20 | 1 | 4 | 1 | - | - | - | - | 1 | - | - | - | - | 1 | - | - |
| **Lafayette** | **15** | 5 | 3 | 7 | 5 | 1 | 4 | 1 | - | - | - | - | - | - | 3 | - | - | 1 | - | - |
| **Lincoln** | **9** | 3 | 3 | 5 | 4 | - | 1 | - | 2 | - | - | - | - | - | - | - | - | - | - | - |
| **Lee** | **8** | 4 | 6 | 3 | 1 | - | 1 | - | - | 1 | - | - | - | - | - | - | - | - | - | - |
| **Lonoke** | **13** | 6 | 4 | 5 | 7 | 3 | 1 | - | - | - | - | - | - | - | - | - | - | - | - | - |
| **Logan** | **41** | 12 | 15 | 3 | 26 | 12 | 13 | - | 1 | - | - | - | - | - | - | - | - | - | - | - |
| **Little River** | **10** | 1 | 4 | 5 | 8 | 1 | 1 | - | - | - | - | - | - | - | - | - | - | - | - | - |
| **Lawrence** | **16** | 1 | 3 | 16 | 2 | 4 | 3 | - | - | - | - | - | - | - | 3 | - | - | - | - | - |
| **Madison** | **36** | 9 | 18 | 24 | 6 | 3 | 6 | 1 | 2 | - | - | - | - | - | 2 | - | - | 1 | - | - |
| *Madison(+)* | *4* | 1 | 1 | 1 | - | 2 | - | 1 | 2 | - | - | - | - | - | - | - | - | - | - | - |
| **Marion** | **40** | 10 | 12 | 11 | 12 | 10 | 7 | - |  | 1 | - | - | 2 | - | 1 | - | - | - | - | - |
| **Miller** | **8** | 3 | 5 | 2 | 1 | 1 | 2 | - | - | - | - | - | - | - | 2 | - | - | - | - | - |
| **Monroe** | **9** | 4 | 1 | 5 | 3 | 4 | 1 | - | - | - | - | - | - | - | - | - | - | - | - | - |
| **Montgomery** | **2** | 1 | 1 | 2 | - | - | - | - | - | - | - | - | - | - | - | - | - | - | - | - |
| **Mississippi** | **2** | 2 | - | 1 | - | - | - | - | 1 | - | - | - | - | - | - | - | - | - | - | - |
| **Nevada** | **14** | 7 | - | 6 | 10 | 2 | 3 | - | - | - | - | - | - | - | - | - | - | - | - | - |
| **Newton** | **216** | 68 | 124 | 49 | 105 | 23 | 48 | - | 5 | 3 | - | 2 | - | - | 2 | - | - | 2 | 1 | - |
| *Newton(+)* | *100* | 28 | 75 | 10 | 54 | 11 | 17 | - | 1 | 1 | - | 1 | - | - | 1 | - | - | - | - | 1 |
| **Ouachita** | **14** | 4 | 3 | 3 | 13 | 3 | 1 | 1 | - | - | - | - | - | - | - | - | - | - | - | - |
| **Perry** | **8** | 4 | 3 | 2 | 2 | - | 2 | 2 | 1 | - | - | - | - | - | - | - | - | - | - | - |
| **Phillips** | **9** | 8 | 1 | 1 | 1 | - | 5 | - | 1 | - | - | - | - | - | - | - | - | - | 1 | - |
| **Pike** | **8** | 2 | 3 | 2 | 5 | - | 2 | 2 | - | - | - | - | - | - | - | - | - | - | - | - |
| **Polk** | **3** | 2 | - | - | - | 2 | 1 | 1 | - | - | - | - | - | - | - | - | - | - | - | - |
| **Poinsett** | **13** | 7 | 1 | 5 | 6 | 2 | 3 | - | 1 | - | - | - | - | - | - | - | - | 1 | - | - |
| **Pope** | **61** | 3 | 48 | 9 | 41 | 3 | 15 | 3 | - | - | - | - | - | - | - | - | - | - | - | - |
| *Pope(+)* | *1* | - | 1 | - | - | - | 1 | - | - | - | - | - | - | - | - | - | - | - | - | - |
| **Prairie** | **11** | 3 | 5 | 7 | 3 | - | 1 | 1 | - | 2 | - | - | - | - | - | - | - | - | - | - |
| **Pulaski** | **11** | 3 | 3 | 4 | 5 | 3 | 3 | 1 | - | - | - | - | - | - | - | - | - | - | - | - |
| **Randolph** | **8** | 3 | 3 | 5 | 3 | - | - | - | 1 | - | - | - | - | - | 1 | - | - | - | - | - |
| **Saline** | **12** | 4 | 4 | 4 | 1 | 3 | 3 | 1 | 2 | - | - | - | - | - | 2 | - | - | - | - | - |
| **Sebastian** | **16** | 6 | 3 | 2 | 12 | 3 | 5 | 1 | - | - | - | - | - | - | - | - | - | - | - | - |
| *Sebastian(+)* | *1* | - | 1 | - | 1 | - | - | - | - | - | - | - | - | - | - | - | - | - | - | - |
| **Scott** | **2** | 1 | 2 | - | 1 | - | - | - | - | - | - | - | - | - | - | - | - | - | - | - |
| **Searcy** | **42** | 8 | 15 | 7 | 31 | 5 | 16 | - | 1 | - | - | - | 1 | - | - | - | - | - | - | - |
| **St Francis** | **8** | 4 | 4 | 3 | 5 | - | - | - | - | - | - | - | - | - | - | - | - | - | - | - |
| **Sharp** | **15** | 3 | 6 | 7 | 8 | 1 | 1 | - | 1 | - | - | - | - | - | 3 | - | - | - | - | - |
| **Stone** | **15** | 6 | 5 | 3 | 6 | 1 | 9 | - | - | - | - | - | - | - | - | - | - | - | - | - |
| **Sevier** | **10** | 5 | 1 | 2 | 5 | 1 | 4 | 2 | - | - | - | - | - | - | - | - | - | - | - | - |
| **Union** | **7** | 3 | 3 | 1 | 4 | 1 | 1 | - | - | - | - | - | - | - | - | - | - | 1 | - | - |
| **VanBuren** | **33** | 7 | 10 | 2 | 21 | 8 | 14 | 2 | - | - | 1 | - | - | - | - | 1 | - | - | - | - |
| **Washington** | **18** | 7 | 5 | 11 | 9 | - | 2 | 1 | 1 | - | - | - | - | - | - | - | - | - | - | - |
| **Woodruff** | **9** | 1 | 2 | 7 | 2 | 1 | 4 | - | - | - | - | - | - | - | - | - | - | 1 | - | - |
| **White** | **8** | 2 | 4 | 6 | - | 1 | 2 | - | 1 | - | - | - | - | - | - | - | - | - | - | - |
| **Yell** | **40** | 9 | 22 | 5 | 21 | 10 | 9 | 3 | 1 | - | - | - | - | - | - | - | - | - | - | - |
| **TOTAL** | **1433** | **435** | **657** | **426** | **657** | **187** | **309** | **42** | **65** | **10** | **2** | **3** | **10** | **2** | **41** | **1** | **1** | **15** | **2** | **1** |

**Supplemental Table 2:** Association of *PRNP* haplotype frequencies and odds ratio with CWD status for Newton County, Arkansas. Haplotypes were derived from unphased sequences of 720 nucleotides of the *PRNP* gene. Haplotype as indicated by letters were also reported by Brandt et al. (2015, 2018), whereas AR_# indicates a haplotype unique to Arkansas. Listed are total numbers (N), relative frequency f(%), and values for deer that tested either CWD-negative (-), CWD-positive (+) or were untested (?). Odds Ratio (OR) reflects relative representation of a haplotype in CWD+ deer, with OR>1 = over-representation and OR<1 = under-representation; SE= standard error, CI= 95% confidence interval, Z(OR)= Z-score and p(OR)= probability. **Values in bold are significant**.

| Hap | Counts |  |  | Frequency (%) |  |  | Odds Ratio |  |  |  |  |
| --- | --- | --- | --- | --- | --- | --- | --- | --- | --- | --- | --- |
|  | N | N(-) | N(+) | f% | f%(-) | f%(+) | OR | SE | CI | Z(OR) | p(OR) |
| A | 95 | 68 | 27 | 15.1 | 15.7 | 13.8 | 0.86 | 0.25 | [0.53-1.38] | -0.64 | 0.52 |
| B | 197 | 124 | 73 | 31.4 | 28.7 | 37.2 | **1.47** | **0.18** | **[1.03-2.11]** | **2.13** | **0.03** |
| C | 59 | 49 | 10 | 9.39 | 11.3 | 5.1 | **0.42** | **0.36** | **[0.21-0.85]** | **-2.42** | **0.02** |
| D | 158 | 105 | 53 | 25.2 | 24.3 | 27 | 1.15 | 0.20 | [0.79-1.70] | 0.73 | 0.46 |
| E | 34 | 23 | 11 | 5.41 | 5.32 | 5.61 | 1.06 | 0.38 | [0.50-2.21] | 0.15 | 0.88 |
| G | 65 | 48 | 17 | 10.4 | 11.1 | 8.67 | 0.76 | 0.30 | [0.43-1.35] | -0.93 | 0.35 |
| I | - | - | - | - | - | - | - | - | - | - | - |
| J | 6 | 5 | 1 | 0.96 | 1.16 | 0.51 | 0.44 | 1.10 | [0.05-3.77] | -0.75 | 0.45 |
| K | 4 | 3 | 1 | 0.64 | 0.69 | 0.51 | 0.73 | 1.16 | [0.08-7.09] | -0.27 | 0.79 |
| L | - | - | - | - | - | - | - | - | - | - | - |
| O | 3 | 2 | 1 | 0.48 | 0.46 | 0.51 | 1.10 | 1.23 | [0.10-12.23] | 0.08 | 0.94 |
| P | - | - | - | - | - | - | - | - | - | - | - |
| R | - | - | - | - | - | - | - | - | - | - | - |
| T | 3 | 2 | 1 | 0.48 | 0.46 | 0.51 | 1.10 | 1.23 | [0.10-12.23] | 0.08 | 0.94 |
| V | - | - | - | - | - | - | - | - | - | - | - |
| AR_1 | - | - | - | - | - | - | - | - | - | - | - |
| AR_2 | 2 | 2 | - | 0.32 | 0.46 | - | - | - | - | - | - |
| AR_3 | 1 | 1 | - | 0.16 | 0.23 | - | - | - | - | - | - |
| AR_4 | 1 | - | 1 | 0.16 | - | 0.51 | - | - | - | - | - |
| Total | 628 | 432 | 196 |  |  |  |  |  |  |  |  |

**Supplemental Table 3:**
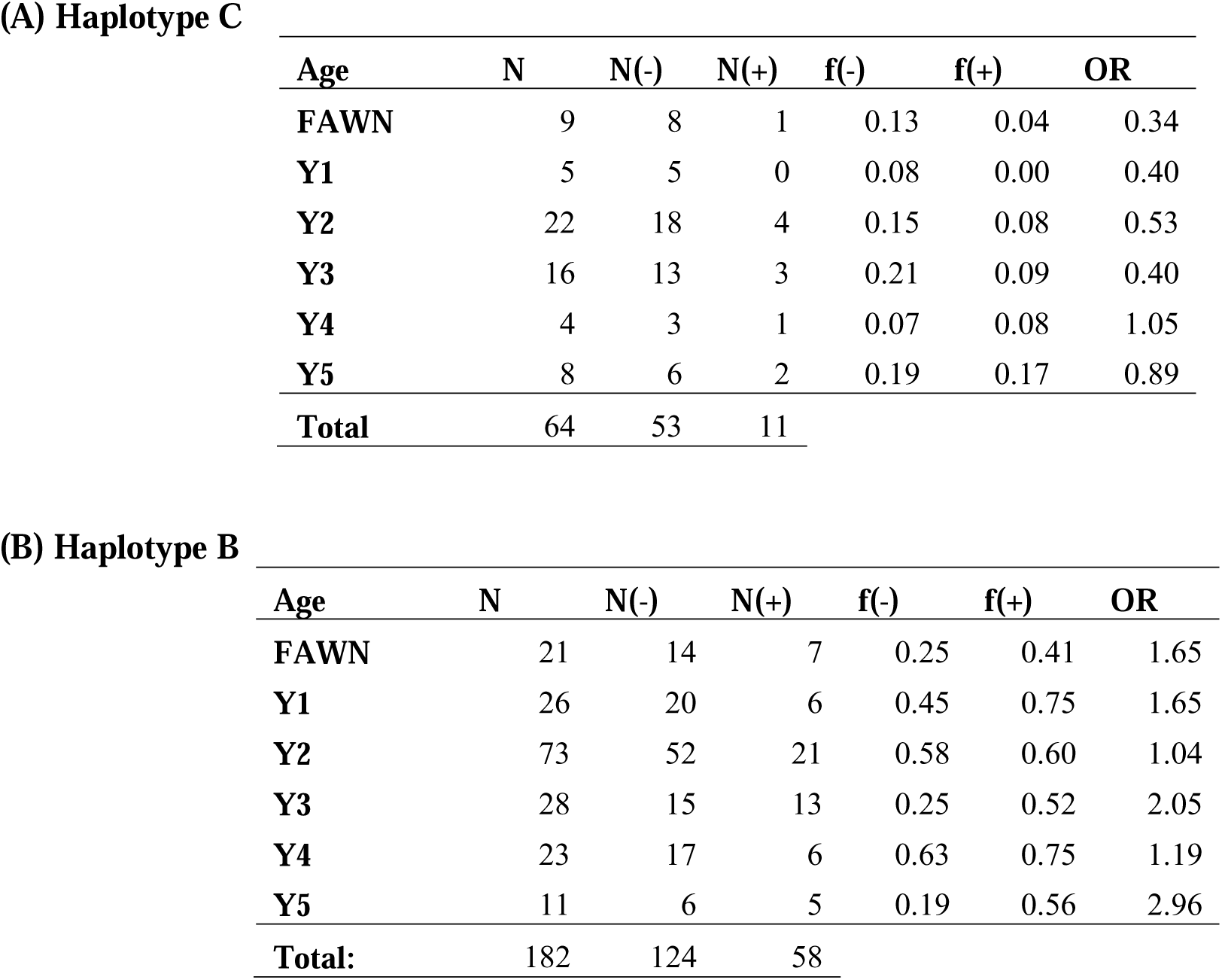
*PRNP* haplotype frequencies and odds ratio for six age classes of white-tailed deer collected in Newton County, Arkansas from 2016-2019. Relative frequencies and odds ratio are shown for (A) Haplotype C (associated with reduced CWD susceptibility), and (B) Haplotype B (associated with increased CWD susceptibility). Relative frequency of CWD-negative [f(-)] and CWD-positive [f(+)] samples were calculated by dividing numbers CWD(+/-) with C or B, respectively, then by negatives w/ other haplotypes.

**Supplemental Table 4:**
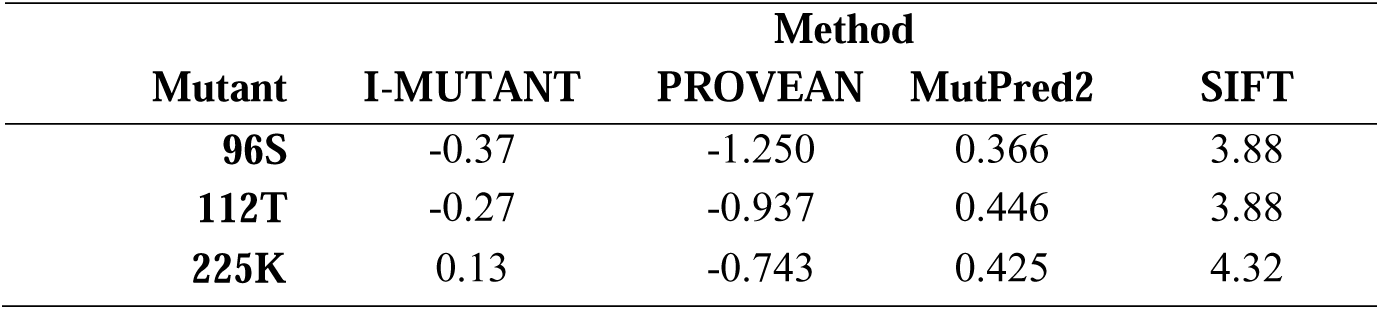
Modeling results for amino acid variant effect versus three non-synonymous *PRNP* gene polymorphisms. Results represent: I-MUTANT stability index (where negative indices destabilize); PROVEAN conservation score (heightened negative value = more likely to be deleterious, with -2.5 as a putative neutral cutoff); MutPred2 probability of pathogenicity (with values >0.8 interpreted as pathogenic); and SIFT conservation score (where 4.32=completely conserved, and 0.0=completely polymorphic).

| Mutant | I-MUTANT | Method |  |  |
| --- | --- | --- | --- | --- |
|  |  | PROVEAN | MutPred2 | SIFT |
| **96S** | -0.37 | -1.250 | 0.366 | 3.88 |
| **112T** | -0.27 | -0.937 | 0.446 | 3.88 |
| **225K** | 0.13 | -0.743 | 0.425 | 4.32 |

**Supplemental Figure 1:**
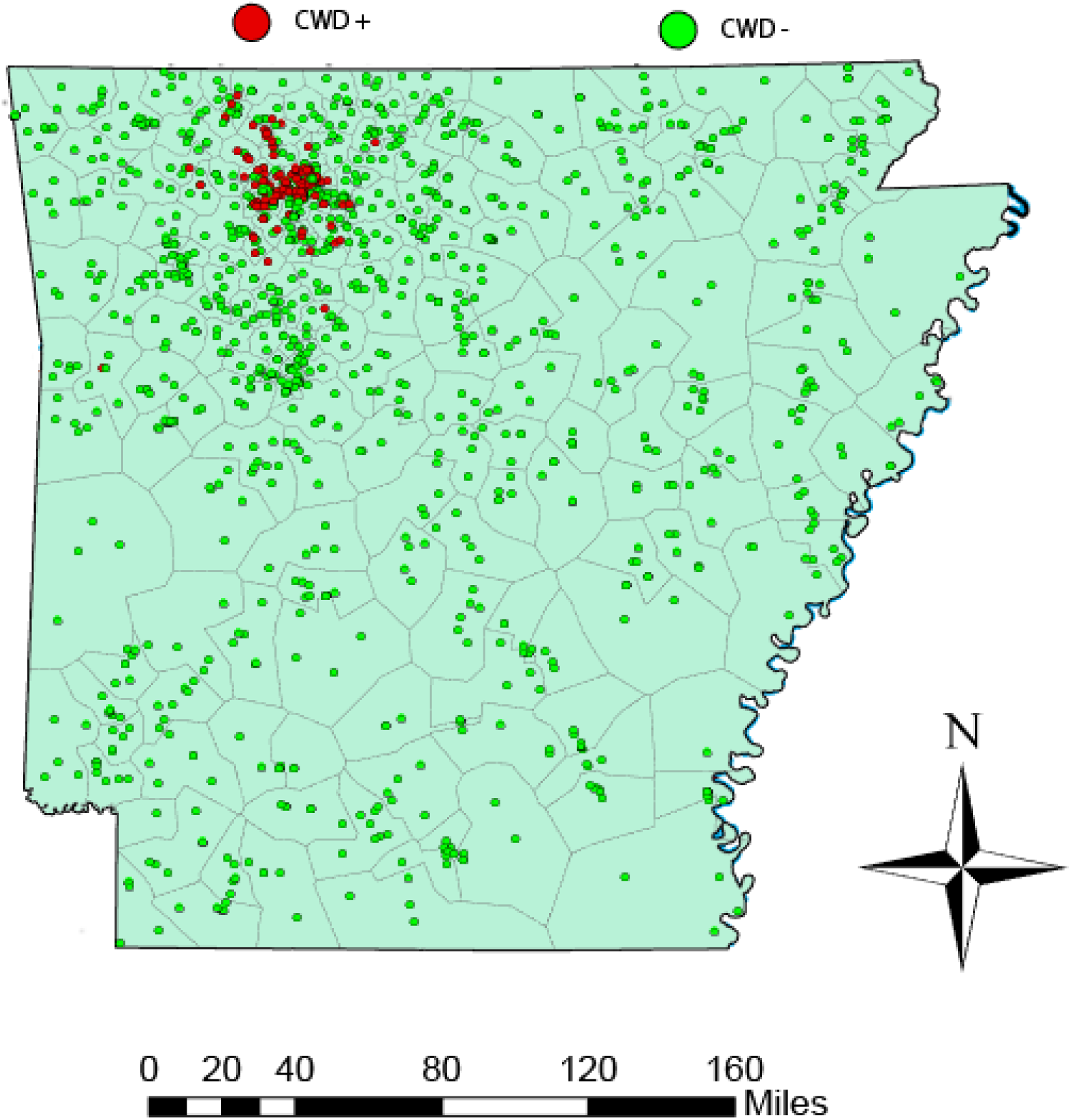
Spatial distribution of 211 non-overlapping polygons in the state of Arkansas (U.S.A.). Each included 5-10 sampling locations. Closed red and green circles represent 1,433 white-tailed deer samples employed to compute prion gene variant (*PRNP* haplotype) frequencies for interpolation. Red circles = CWD+; green circles = CWD-

**Supplemental Figure 2:**
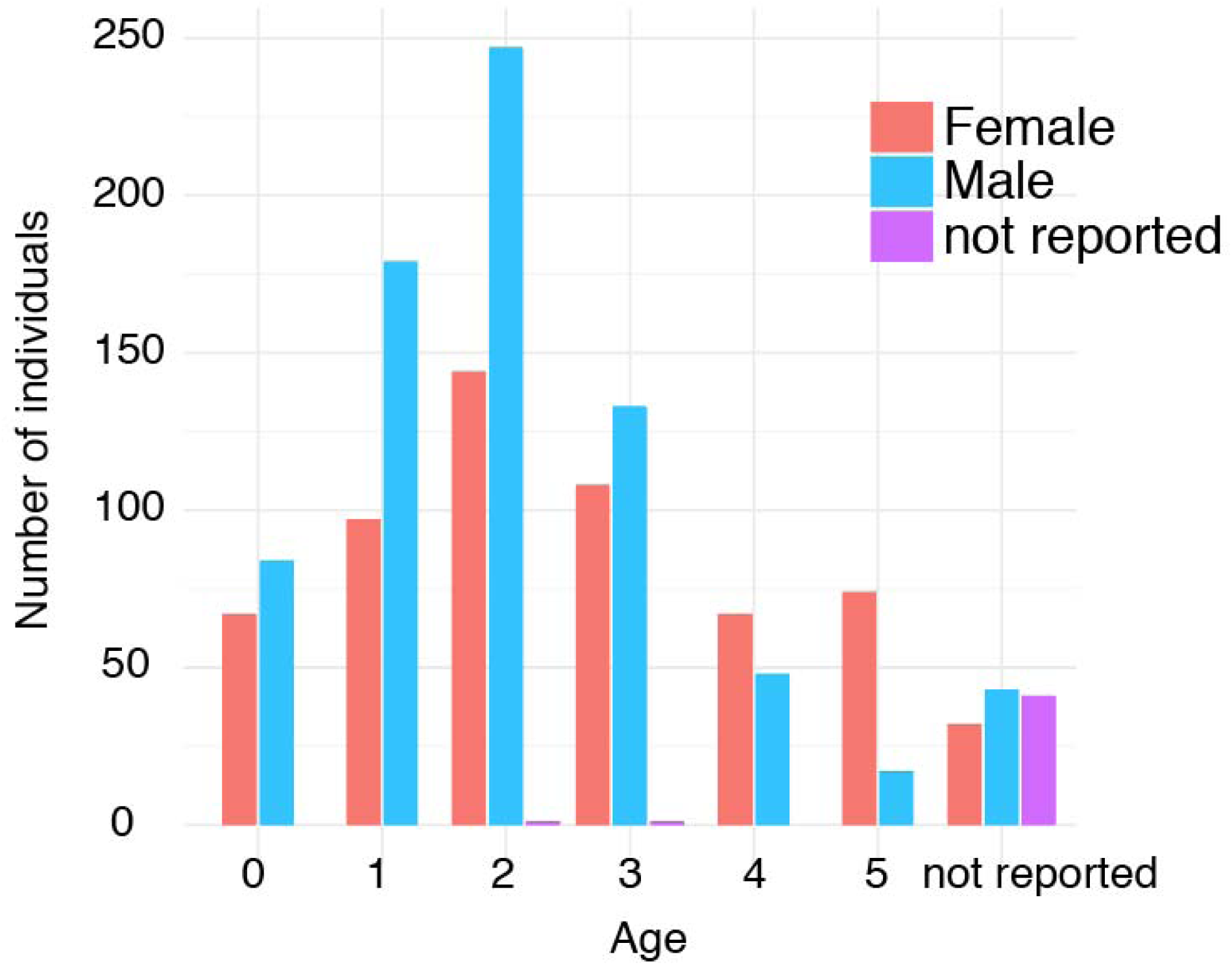
Age and sex distribution for *N*=1,433 white-tailed deer tissue sample for which *PRNP* haplotype data was generated.

## Notes

### Competing Interest Statement

The authors have declared no competing interest.

